# Image-based computational fluid dynamics to compare two mitral valve reparative techniques for the prolapse

**DOI:** 10.1101/2023.12.22.572827

**Authors:** Lorenzo Bennati, Giovanni Puppini, Vincenzo Giambruno, Giovanni Battista Luciani, Christian Vergara

**Affiliations:** Department of Surgery, Dentistry, Pediatrics, and Obstetrics/Gynecology, University of Verona, Piazzale Ludovico Antonio Scuro 10, Verona, 37134, Italy; Department of Radiology, University of Verona, Piazzale Stefani 1, Verona, 37126, Italy; Division of Cardiac Surgery, Department of Surgery, Dentistry, Pediatrics, and Obstetrics/Gynecology, University of Verona, Piazzale Stefani 1, Verona, 37126, Italy; LaBS, Dipartimento di Chimica, Materiali e Ingegneria Chimica “Giulio Natta”, Politecnico di Milano, Piazza Leonardo da Vinci 32, Milan, 20133, Italy

**Author notes:** Contributing authors.

**Keywords:** mitral valve prolapse, neochordae technique, resection technique, computational fluid dynamics, turbulence, hemolysis

## Abstract

**Objective:** Nowadays, the treatment of mitral valve prolapse involves two distinct repair techniques: chordal replacement (Neochordae technique) and leaflet resection (Resection technique). However, there is still a debate in the literature about which is the optimal one. In this context, we performed an image-based computational fluid dynamic study to evaluate blood dynamics in the two surgical techniques.

**Methods:** We considered a healthy subject (H) and two patients (N and R) who underwent surgery for the prolapse of the posterior leaflet and were operated with the Neochordae and Resection technique, respectively. Computational Fluid Dynamics (CFD) was employed with prescribed motion of the entire left heart coming from cine-MRI images, with a Large Eddy Simulation model to describe the transition to turbulence and a resistive method for managing valve dynamics. We created three different virtual scenarios where the operated mitral valves were inserted in the same left heart geometry of the healthy subject to study the differences attributed only to the two techniques.

**Results:** We compared the three scenarios by quantitatively analyzing ventricular velocity patterns and pressures, transition to turbulence, and the ventricle ability to prevent thrombi formation. From these results we found that both the operated cases were able to restore almost physiological blood dynamic conditions, with some differences due to the reduced mobility of the Resection posterior leaflet. Conclusions: Our findings suggest that the Neochordae technique developed a slightly more physiological flow with respect to the Resection technique. The latter gave rise to a different direction of the mitral jet during diastole increasing the turbulence that is associated with ventricular effort and hemolysis, with also a larger ability to washout the ventricular apex preventing from thrombi formation.

## 1. Introduction

Mitral Valve Prolapse (MVP) is a valvular disease characterized by an unphysiological displacement of the leaflets towards the left atrium during the systolic phase due to elongated or broken chordae tendineae. The main consequence is mitral regurgitation.

Nowadays, there are two different surgical reparative techniques to treat MVP: the *Resection* technique and the *Neochordae* technique. In the first one, the surgeon excises the prolapsed portion of the leaflet (and associated ruptured or elongated chordae) and re-approximate the surrounding leftover leaflet tissue, restoring the coaptation surface [1–3]. This technique has been established as the gold standard for the treatment of posterior leaflet MVP [2, 3]. However, the main disadvantage is related to the fact that the higher the portion of tissue removed, the lower the leaflet mobility, resulting, in the worst scenario, in a mono-cuspid mitral valve, see Figure 1, left [4].

**Fig. 1.**
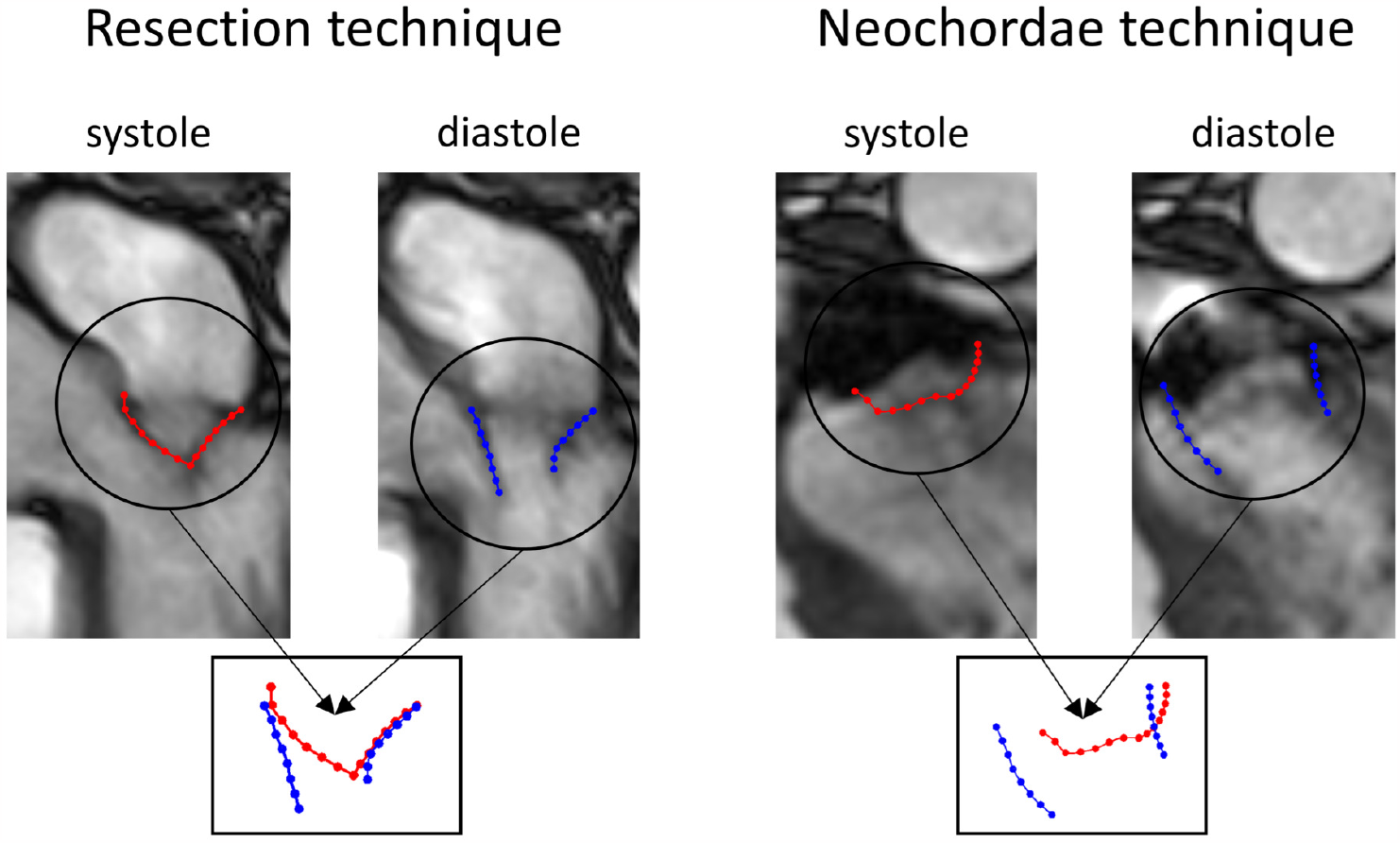
Left: Cine-MRI images of a patient operated with Resection technique to treat the posterior leaflet prolapse of the P2 segment. We reported the leaflets contours in closed (red) and open (blue) configurations. Notice that the posterior (operated) leaflet has a reduced mobility as a consequence of the reparative technique; Right: Cine-MRI images of a patient operated with the Neochordae technique. Notice that this technique ensures greater mobility of the posterior leaflet.

To overcome the hypo-mobility of the operated leaflet, Neochordae technique has been introduced [5, 6]. This method is also called *Respect* approach as repair involves placement of artificial chordae and a minimal to no leaflet resection. In particular, a set of pre-made polytetrafluoroethylene (ePTFE) chordae are anchored to the papillary muscles and then used to resuspend the prolapsed segment of the leaflet, see Figure 1, right. The main drawback of this technique is the determination of the appropriate length for neochordae [7] that often requires a trial-and-error approach during the surgical procedure. Although the long-term results are promising [8], there is still debate in the clinical world about which techniques to use and their effects on ventricular fluid dynamics [3].

In this respect, computational methods represent a valuable and non invasive tool to quantitatively assess the 3D local velocity patterns in the heart chambers and areas of disturbed flow to enhance the understanding of cardiac physio-pathology [9–14] and the design of valve prosthesis and surgical interventions [13, 15–18]. In particular, some works addressed the issue of studying the two surgical techniques mentioned above. This can be grouped in two categories: *Structure-only* (S), where no blood dynamics is simulated, and *Fluid-Structure Interaction* (FSI) models. Regarding the first approach, we cite [19] where the authors developed a computational simulation protocol to perform virtual Resection technique, whereas [20] investigated different neochordae implantation sites and [21, 22] different neochordae tensioning and lengths. At the best of our knowledge, only [23] performed a virtual comparison of the two reparative techniques starting from an ideal mitral valve. For the FSI approach, we cite [13] where the optimal number of neochordae has been investigated in different types of prolapse obtained by virtually deforming a healthy mitral valve.

In this context, the aim of our work is to compare the two reparative techniques to investigate their effect on the ventricular blood flow. To do this we used computational fluid dynamics (CFD) with imposed motion where the displacement of the left heart (left ventricle, left atrium, aortic root, mitral and aortic valve) is provided by dynamic imaging, (*Dynamic Image-Based CFD*, DIB-CFD). The choice of using a DIB-CFD model is motivated by the availability of time-resolved cine-MRI images of two repaired mitral valves together with the left heart wall motion. This allows us to focus on and compare hemodynamic quantities such as velocity, pressure, Wall Shear Stresses (WSS), and turbulence [24–28].

Specifically, we reconstructed three different mitral valves (one healthy and two operated with the two techniques) geometries and motion, and we created three different virtual scenarios where the mitral valves are inserted in the same healthy ventricular geometry and motion, supposed to be the same for the three cases. DIB-CFD is then run for the three cases in the same haemodynamic settings. This virtual comparison is employed to emphasize heamodynamic variations due solely to the geometric differences induced by the different techniques.

To the best of our knowledge, the present study features two novelties:

- The reconstruction of the patient-specific mitral valve geometries of two cases, one for each of the repair surgical techniques. Specifically, we also reconstructed the patient-specific motion of such valves;
- A computational comparison of haemodynamics in the two repair technique configurations, including the potential risk of red blood cell damage, ventricular washout, and remodeling indices [29, 30].

To perform this comparison, we used a fluid dynamic incompressible model for blood with a resistive method to treat the presence of the valves and a Large Eddy Simulation (LES) model to account for the transition to turbulence, and we compared different haemodynamic meaningful indices.

## 2 Materials and Methods

In this section, we first described the procedure to obtain the geometries and motion of the left heart internal wall and valves used to create the three virtual scenarios. Then, we briefly reported the mathematical and numerical methods used in this work. Finally, we introduced the quantities of interest that has been analyzed in the Results section.

### 2.1 Creation of the virtual scenarios

We considered a healthy subject (H) and two patients (N and R) operated at Borgo Trento Hospital in Verona with the Neochordae and Resection techniques, respectively, due to the prolapse of the P2 segment of the posterior leaflet. For each of them, we have at disposal cine-MRI images.

We started by the reconstructed motion of the internal wall of the left ventricle, left atrium, and aortic root of case H, obtained from 30 cine-MRI frames as reported in [31]. In Figure 2A, we display this left heart (LH) displacement 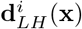, *i* = 1, …, 30, (with respect to the end-systolic ventricular configuration) at three representative frames, together with the corresponding time evolution of ventricular volume and flow rate through aortic and mitral orifices computed by means of the cine-MRI images, see Figure 2B and Figure 2C. Moreover, we have at disposal also the aortic valve (AV) geometry in the closed and open configurations, see Figure 2D. From these we computed the displacement **d**_*AV*_ (**x**) as the difference between the open and closed states.

**Fig. 2.**
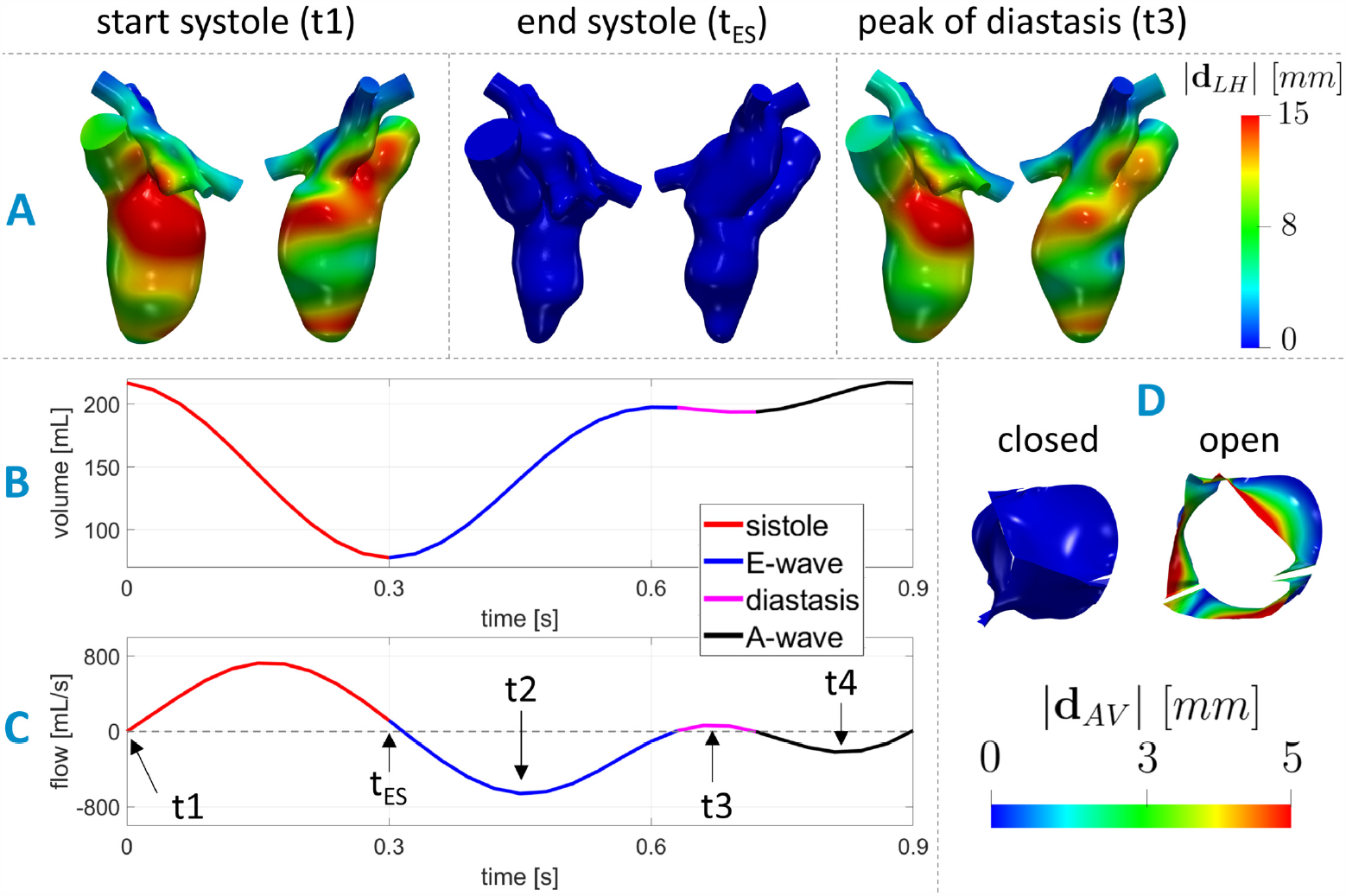
Geometric and dynamic data taken from [24]. A: Two views of the geometries and magnitude of the LH displacement 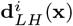, *i* = 1, …, 30, at the begin of systole t1 (closed mitral valve), end systole (*t*_*ES*_) (closed mitral valve), the instant t2 of maximum valve opening (peak of flow rate during E-wave), the instant t3 of maximum ventricular flow rate during diastasis (i.e. the period of partial closure between the two waves), and the instant t4 of maximum valve opening during the A-wave (peak of flow rate during A-wave); B: Trend in time of the ventricular reconstructed volume; C: Flow rate through the aortic (from 0 to 0.3 s) and mitral orifice (from 0.3 to 0.9 s). The arrows refer to the end-diastolic frame (t1), end-systolic frame (*t*_*ES*_), peak of E-wave (t2), peak of diastasis (t3) and peak of A-wave (t4); D: Geometries and magnitude of the aortic valve displacement **d**_*AV*_ (**x**) in its fully closed and fully open configuration.

For case H we have also at disposal cine-MRI images of the mitral valve consisting in 30 frames per heartbeat acquired following the *ad hoc* protocol proposed in [32] and based on a radial sampling, which are here reconstructed for the first time. Specifically, we reconstructed the geometries and the displacement 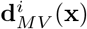, *i* = 1, …, 30, at all the frames, employing the method proposed in [24, 32] (see the Results Section). Furthermore, for each of the two operated cases (N and R), new cine-MRI images of the mitral valve, consisting in 30 acquisitions per heartbeat, were provided. Ethical review board approval and informed consent were obtained from both the patients. The image acquisitions were performed few days after the intervention. Also for these operated cases, we reconstructed the geometries and the displacement 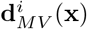 of the mitral valve at all the available frames (see the Results Section). Then, the geometries of the two mitral valves were virtually inserted and adapted to the 30 reconstructions at disposal of the left heart of the healthy subject H, see Figure 2A. This was done by using the same ratio between the area of the annulus and the area of the mitral orifice at the peak of the E-wave measured for the patients from imaging. In particular, we have 0.52 and 0.63 for N and R, respectively.

We remind that the idea of this work is to compare, by means of a computational analysis, the blood dynamics in the two post-operative scenarios N and R with the healthy case, by using for all the three scenarios the same ventricular geometry and motion, thus highlighting the differences due to only the reparative technique. In Figure 3, we displayed the three reconstructed virtual scenarios.

**Fig. 3.**
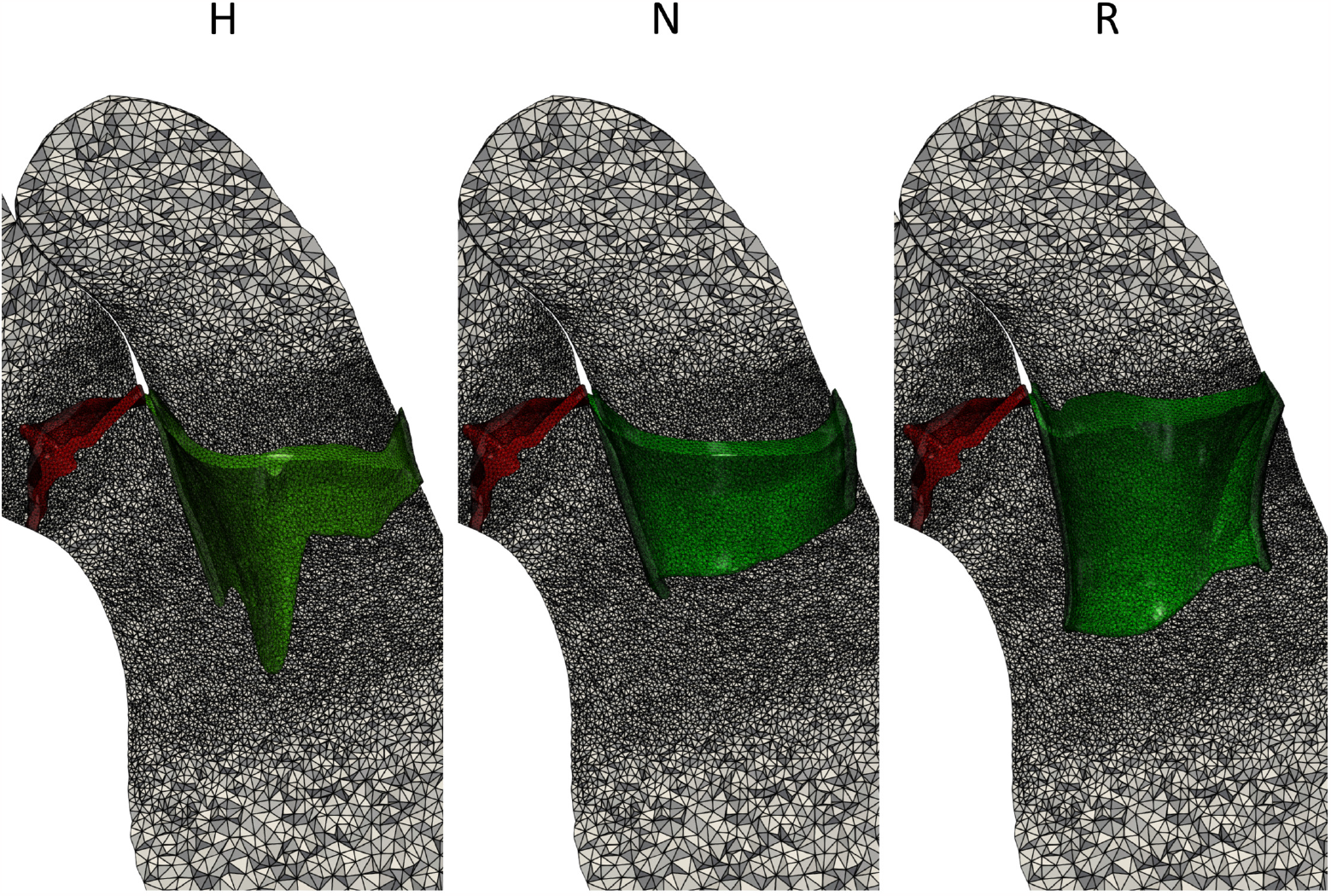
Computational mesh of three virtual scenarios: Healthy (H), Neochordae (N) and Resection (R). In red we displayed the aortic valve and in green the mitral valve.

### 2.2 Mathematical and numerical modeling

We considered blood as an incompressible, homogeneous, Newtonian fluid with density *ρ* = 1.06 · 10^3^ *kg/m*^3^ and dynamic viscosity *μ* = 3.5 · 10^*−*3^ *Pa · s*, described by the Navier-Stokes (NS) equations, see [33, 34]. To solve NS in the moving LH we used the Arbitrary Lagrangian Eulerian (ALE) framework [35] and to manage the presence of the valves we used the Resistive Immersed Implicit Surface (RIIS) method [36, 37]. To evaluate the transition to turbulence occurring in the left heart [38], we employed the *σ*-LES method proposed for ventricular blood dynamics in [39] and successfully used in different hemodynamic applications [24, 40–42]. See [31] for further details. In particular, the displacement of LH 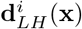 is derived in time and used to compute the wall velocity to prescribe it as boundary condition for the NS equations. However, 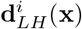 has been obtained only at the 30 MRI acquisition times, thus we performed a spline interpolation to obtain **d**_*LH*_ (**x**, *t*) for all *t ∈* [0, *T*] where *T* = 0.9 *s* is the duration of the heartbeat. According to the ALE framework, at each time, the fluid domain Ω(*t*) is obtained by extending **d**_*LH*_ (**x**, *t*) into Ω through the solution of a linear elastic problem [43].

In Figure 4A, we displayed the fluid domain, where Σ_*LH*_ represents the internal wall surfaces of LH, Σ_*AR*_ and Σ_*P V*_, the outlet and inlet sections of the aortic root and pulmonary veins, respectively. In yellow we reported the aortic valve Γ_*AV*_ and in green the mitral valve Γ_*MV*_ of the healthy subject. Thus, the ALE NS equations in the known domain Ω(*t*) are solved to find the blood pressure *p* and the blood velocity **u**:

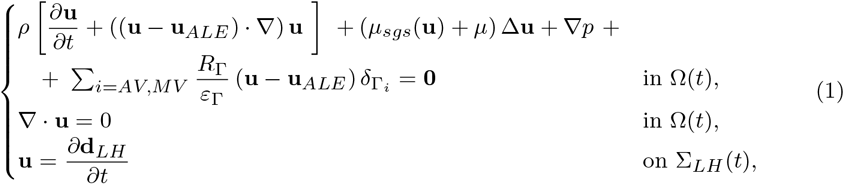

with a null initial condition in Ω(0). *μ*_*sgs*_(**u**) is the sub-grid viscosity of the *σ*-model [39]; 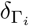 is a smoothed Dirac delta function representing a layer, with thickness 2*ε*_Γ_, around the surface of the valve Γ_*i*_, *i* = *AV, MV*, [11, 37]; *R*_Γ_ is a penalization term used to enforce the kinematic constraint. In our numerical simulations, we set *R*_Γ_ = 10^5^ *kg/m · s* and *ε*_Γ_ = 0.75 *mm* [14, 24, 31, 44, 45]. The position of the surface of each valve Γ_*i*_, *i* = *AV, MV*, is updated at each time according to the valves displacement. In particular, for the AV case, since the displacement has been defined only in the open and closed configurations (see Figure 2D), we multiplied **d**_*AV*_ (**x**) by a linear coefficient *C*_*AV*_ (*t*), *t ∈* [0, *T*], equal to 0 when the valve is closed and 1 when it is opened. The opening and closing duration has been set according to the literature [46], i.e. 19 and 47 ms, for the opening and closing, respectively. Conversely, for MV we did not need to make assumptions on the opening and closing times, since we reconstructed all the configurations from cine-MRI images. Thus, 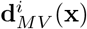 has been directly interpolated in time to obtain **d**_*MV*_ (**x**, *t*), *t ∈* [0, *T*].

**Fig. 4.**
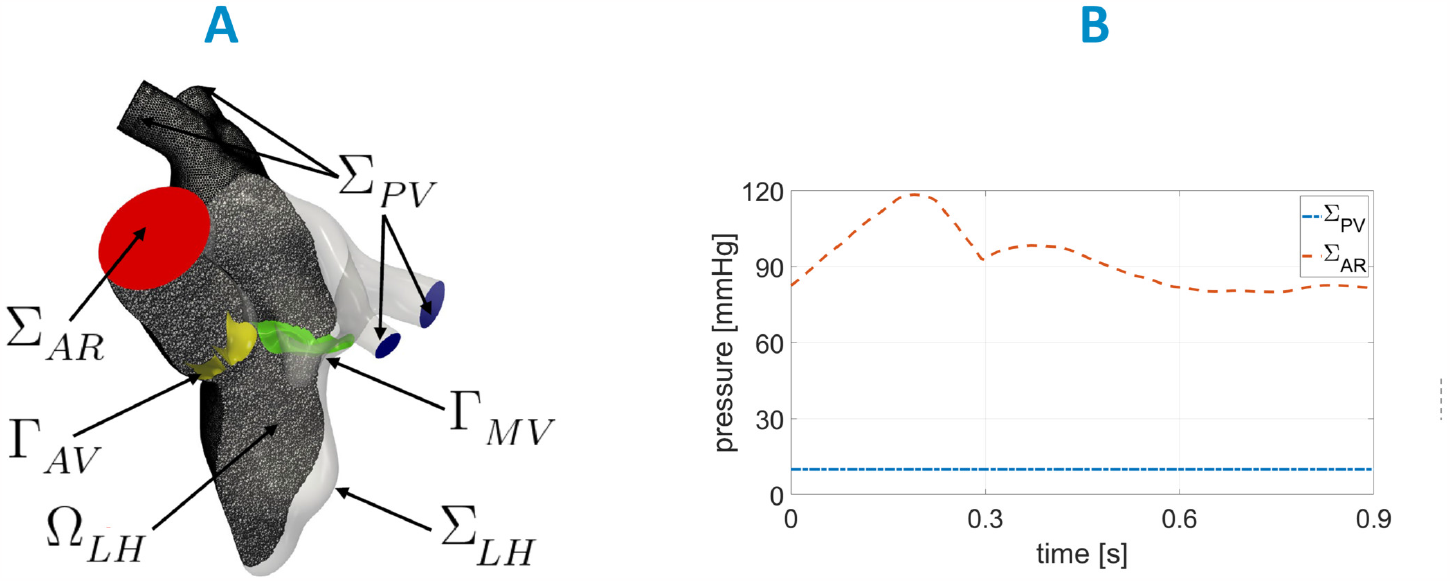
A: Computational domain Ω with its boundaries. In yellow we reported the aortic valve Γ_*AV*_ and in green the mitral valve Γ_*MV*_ of H. B: Trend in time of the pressures imposed at Σ_*P V*_ and Σ_*AR*_ for the three scenarios.

Regarding the remaining boundary conditions of system (1), for all the tree cases we prescribed a constant pressure of 10 *mmHg* on Σ_*P V*_ [47, 48] (Neumann condition on the normal direction), and a time dependent physiological pressure taken from the Wiggers diagram [47, 49] at Σ_*AR*_, see Figure 4B. In the tangential direction, in order to avoid possible backflows instabilities, we prescribed a null velocity both on Σ_*P V*_ and Σ_*AR*_ [50].

To numerically solve system (1), we used first-order Finite Elements together with first order semi-implicit discretization in time [51]. The numerical scheme was stabilized by means of the SUPG-PSPG scheme [52] implemented in the multiphysics high performance library *life*^*x*^ [53, 54] (https://lifex.gitlab.io/) based on the deal.II core [55]. We run the simulations using 192 parallel processes on the GALILEO100 supercomputer (https://www.hpc.cineca.it/hardware/galileo100) at the CINECA high-performance computing center (Italy).

Tetrahedral mesh of the left heart was generated in Vascular Modeling ToolKit (VMTK) [56, 57] with an average mesh element size of 0.9 mm and a local refinement in correspondence of the valves of 0.3 mm, see Figure 3. The timestep Δ*t* was equal to 5·10^*−*4^ *s*. We performed a mesh convergence test ensuring that no significant differences may be found by using a finer mesh or a smaller timestep. Furthermore, with this value of the average mesh element size, we are able to satisfy the Pope criterion used to assess the LES quality [58]. In particular, we computed the quantity *M* (**x**, *t*):

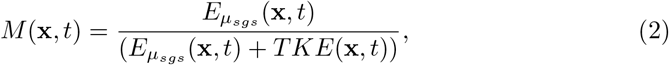

where 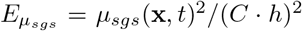 [59] is the turbulent kinetic energy related to the unresolved scales, where *C* = 1.5 is the LES constant [39] and *h* is the local cell diameter; *TKE* is the turbulent kinetic energy of the resolved scales. Values of *M* below the threshold of 20% indicate that the LES is sufficiently resolved [26, 38, 58]. In our simulations, the average in time of the left ventricle volume with M below this threshold was about 80% for all the three scenarios, confirming that with such value of the average mesh element size we were able to capture, on average, 80% of the turbulent kinetic energy of the left ventricle. This result is in accordance with that found in other ventricular LES studies, see, e.g, [24, 26, 38].

### 2.3 Quantities of interest

We computed the ensemble velocity (i.e. the average calculated over 9 heartbeats) and to compare and quantify the effects of the two repair techniques on the ventricular flow we introduced the following post-processed quantities:

- *Flow Stasis (FS)*: is a function of space representing the percentage of the heartbeat during which the velocity magnitude is smaller than 0.1 m/s as suggested in [60–62]. High values of FS in correspondence of the ventricular apex may indicate a low washing out ability that may promote the thrombi formation;
- *Turbulent Kinetic Energy (TKE)*: at each time and space quantifies the velocity fluctuations by means of the fluid Reynolds stress tensor [26, 38]. High values of TKE in the ventricle may be correlated with a non-physiological increased ventricular effort [26, 30];
- *Vorticity* : at each time and space quantifies the amount of rotational behaviour of blood flow [26, 38]. Together with turbulence, it allows to quantify the disturbed flow developed in the heart chambers;
- *Turbulent force τ*_*max*_ (obtained from the fluid Reynolds stress tensor [63]): is a function of space and time quantifying the turbulent forces exerted by the fluid on the red blood cells. Values exceeding 800 Pa are recognized as condition that can damage the red blood cells promoting hemolysis [63].

## 3 Results

In Figure 5, we reported the geometries and the displacement magnitude 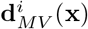 of the reconstructed mitral valves of H, N and R, at four representative instants. The values of the displacement were calculated with respect to the initial systolic configuration at t1. We noticed that at t2 and t4 all three scenarios featured a comparable orifice area, whereas at the peak of diastasis t3, R had a smaller area due to a less posterior leaflet mobility.

**Fig. 5.**
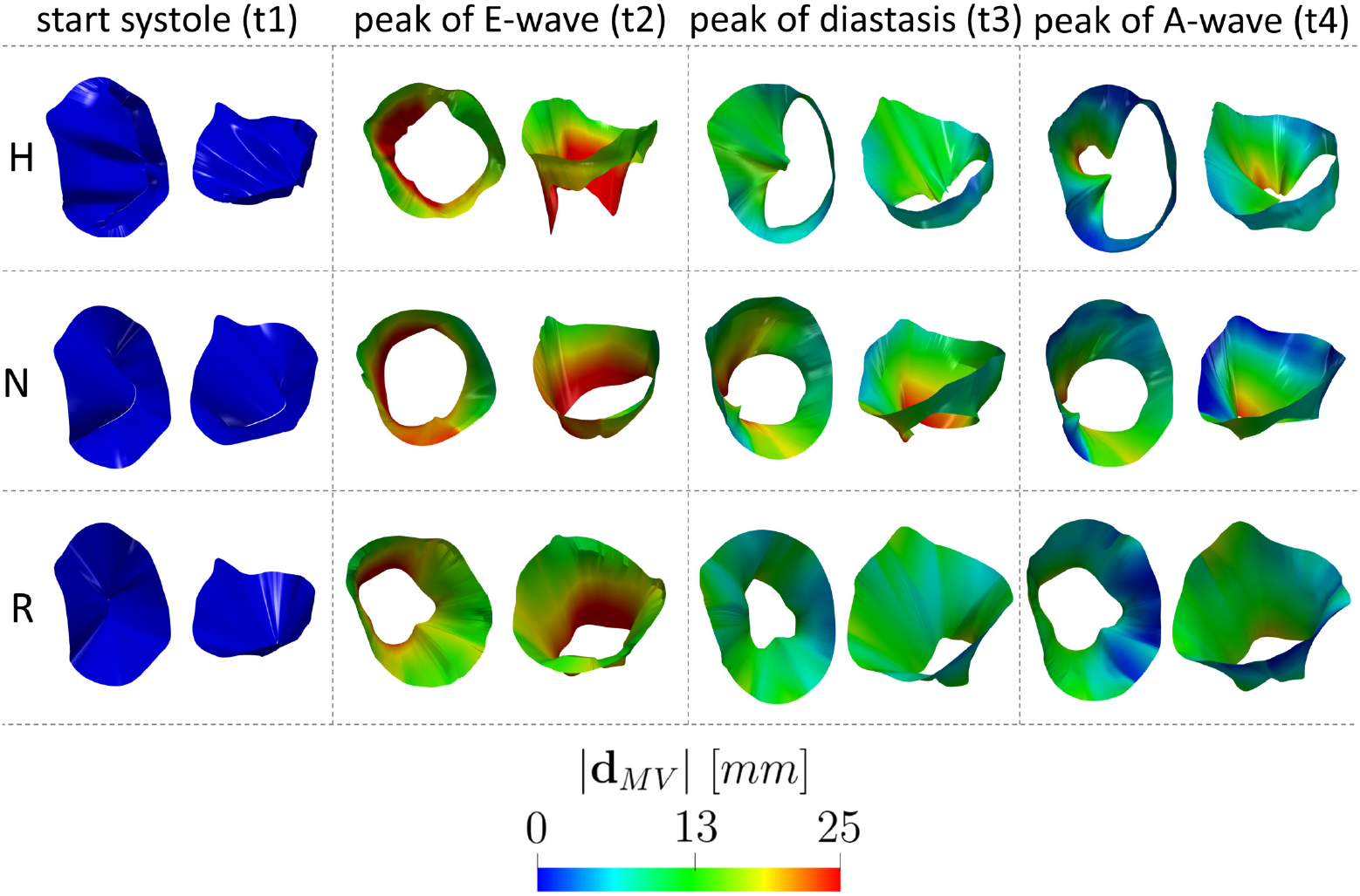
Geometries and magnitude of the reconstructed cine-MRI displacement 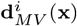, *i* = 1 …, 30, of the three mitral valve configurations at (see Figure 2C): the begin of systole t1, peak of flow rate during E-wave t2, the instant t3 of maximum ventricular flow rate during diastasis, and the instant t4 of maximum valve opening during the A-wave, for the three scenarios Healthy (H), Neochordae (N), and Resection (R). The displacement was calculated with respect the start-systolic configuration.

In Figure 6, we reported a longitudinal slice with the spatial distribution of the ensemble velocity magnitude at three instants for each scenario. At the peak of the E-wave t2, we observed comparable velocities across the mitral valve in all the scenarios with the formation of two ventricular vortex rings below the leaflets. However, in R the mitral jet is oriented more towards the apex, while in scenarios H and N, the jet develops more along the ventricular wall. Additionally, we noticed in N and R the formation of a clockwise vortex in correspondence of the apex. At the peak of diastasis t3, we noticed a uniform clockwise vortex in H and N in the middle of the ventricle, whereas in R more swirling and chaotic structures were present. At the peak of A-wave t4, during the second injection of fluid in the ventricle, the velocities through MV in all the scenarios were too low to reach the middle of the ventricle, where the vortexes formed during diastasis were still present.

**Fig. 6.**
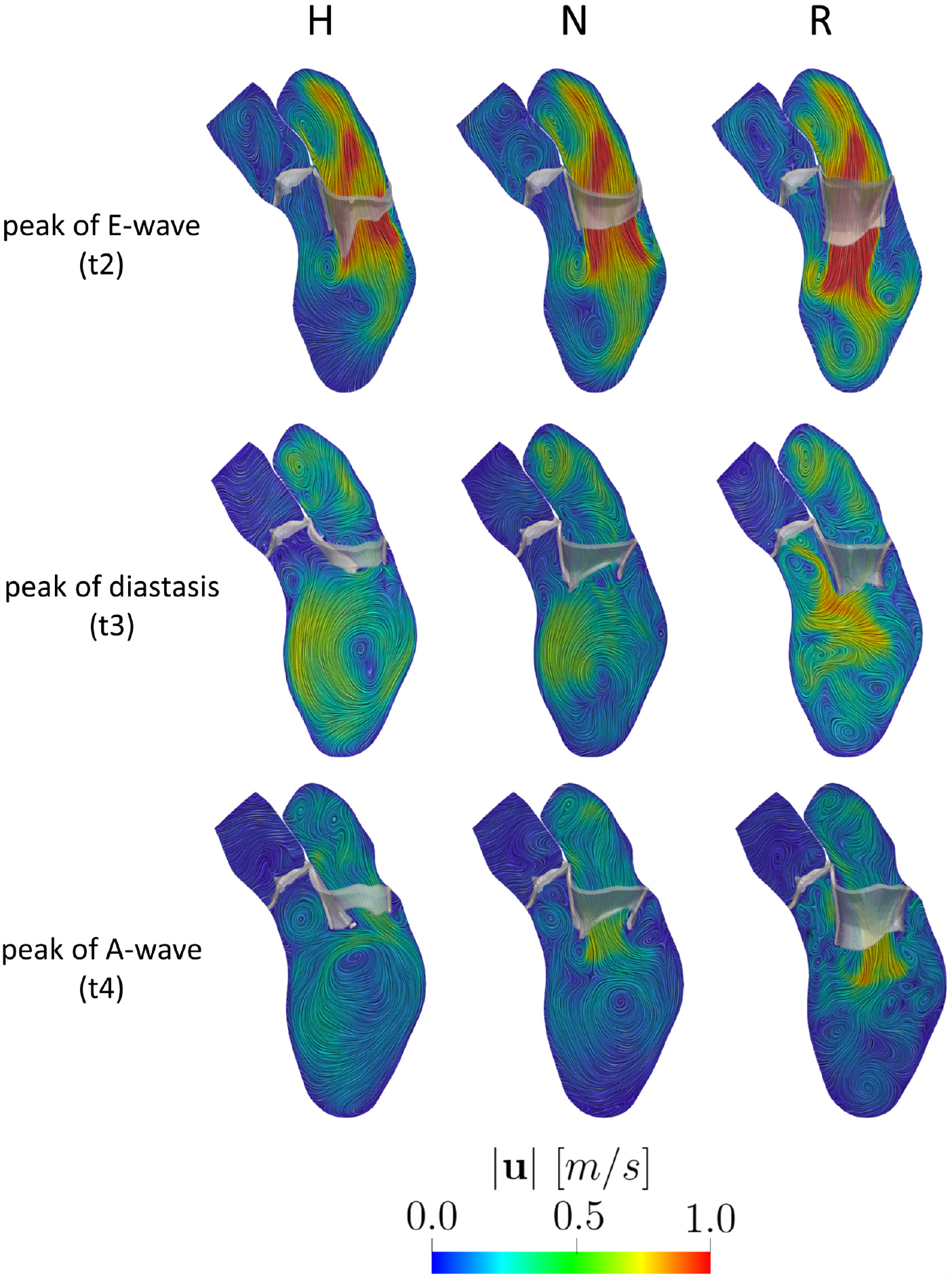
Magnitude of the ensemble velocity computed over 9 heartbeats at t2 (peak E-wave), t3 (peak of diastasis), and t4 (peak of A-wave) for the three scenarios Healthy (H), Neochordae (N) and Resection (R).

In Figure 7, we reported the spatial distribution of the ensemble pressure together with the ensemble velocity patterns at the peak of the E-wave for the three scenarios. For H and N, the pressure appears to be the same and homogeneous in the atrium and in the ventricle, whereas in R a negative pressure (reaching values up to -5 mmHg) is developing in the middle of the ventricle due to the presence of more pronounced eddies below the MV leaflets (cfr Figure 6).

**Fig. 7.**
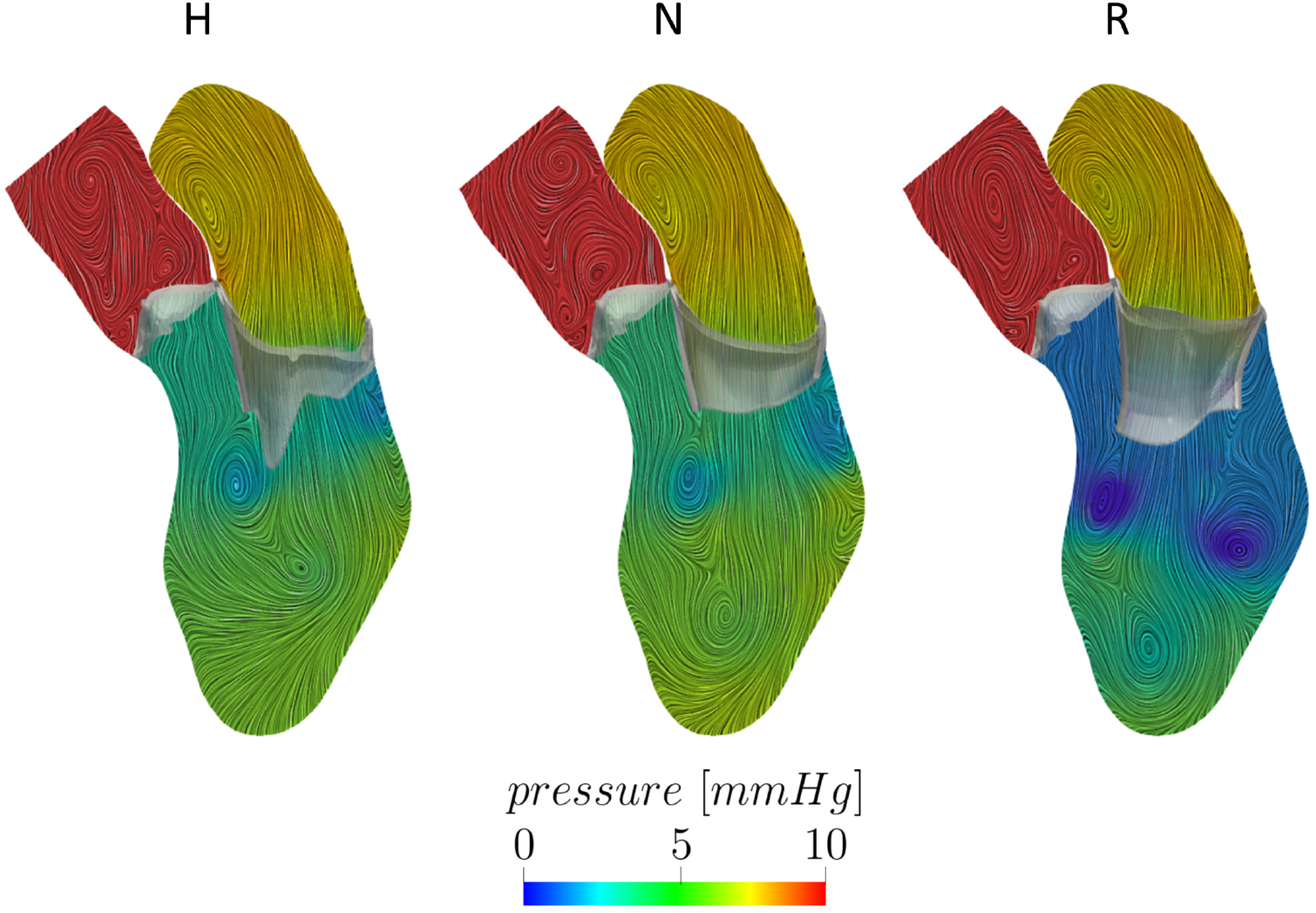
Spatial distribution of the ensemble pressure computed over 9 heartbeats with the ensemble velocity patterns in background at the peak of the E-wave t2 for the three scenarios Healthy (H), Neochordae (N) and Resection (R)

In Figure 8, top, we reported the pressure drops evaluated between spheres placed in the transversal and longitudinal (trans-valvular) directions. The first one is significant to analyze possible remodeling of the ventricle [30], whereas the second is standard clinical measure to assess the functioning of the mitral valve. In both the cases we noticed that H and N exhibited a comparable pattern, whereas R displayed slightly larger oscillating values during the E-wave. This is particularly evident for the trans-valvular pressure drop, see also Figure 7. In Table 1, for each scenario we reported the average in time (during diastole) of the two pressure drops. We observed results very similar to the healthy case for both the repair techniques in the transversal direction, whereas in the trans-valvular direction the Resection technique exhibits larger values if compared with the healthy case and the Neochordae technique.

**Table 1.**
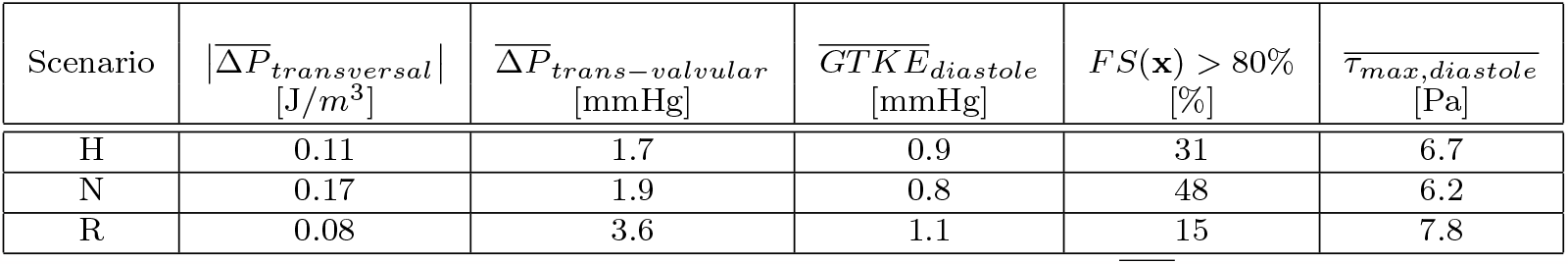
Values of the quantities of interest computed for each scenario. 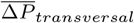: average in time of the pressure drop evaluated between the ventricular septum and free wall during the diastolic phase; 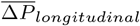 average in time of the pressure drop evaluated between the ventricular base and apex during the diastolic phase; 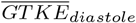: average in time of GTKE evaluated in the ventricle during the diastolic phase; Percentage of volume with FS(**x**) greater than 80% evaluated in the ventricular apex; 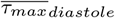: average in time of *τ*_*max*_ evaluated in the ventricle during the diastolic phase. Scenarios: Healthy (H), Neochordae (N), and Resection (R).

**Fig. 8.**
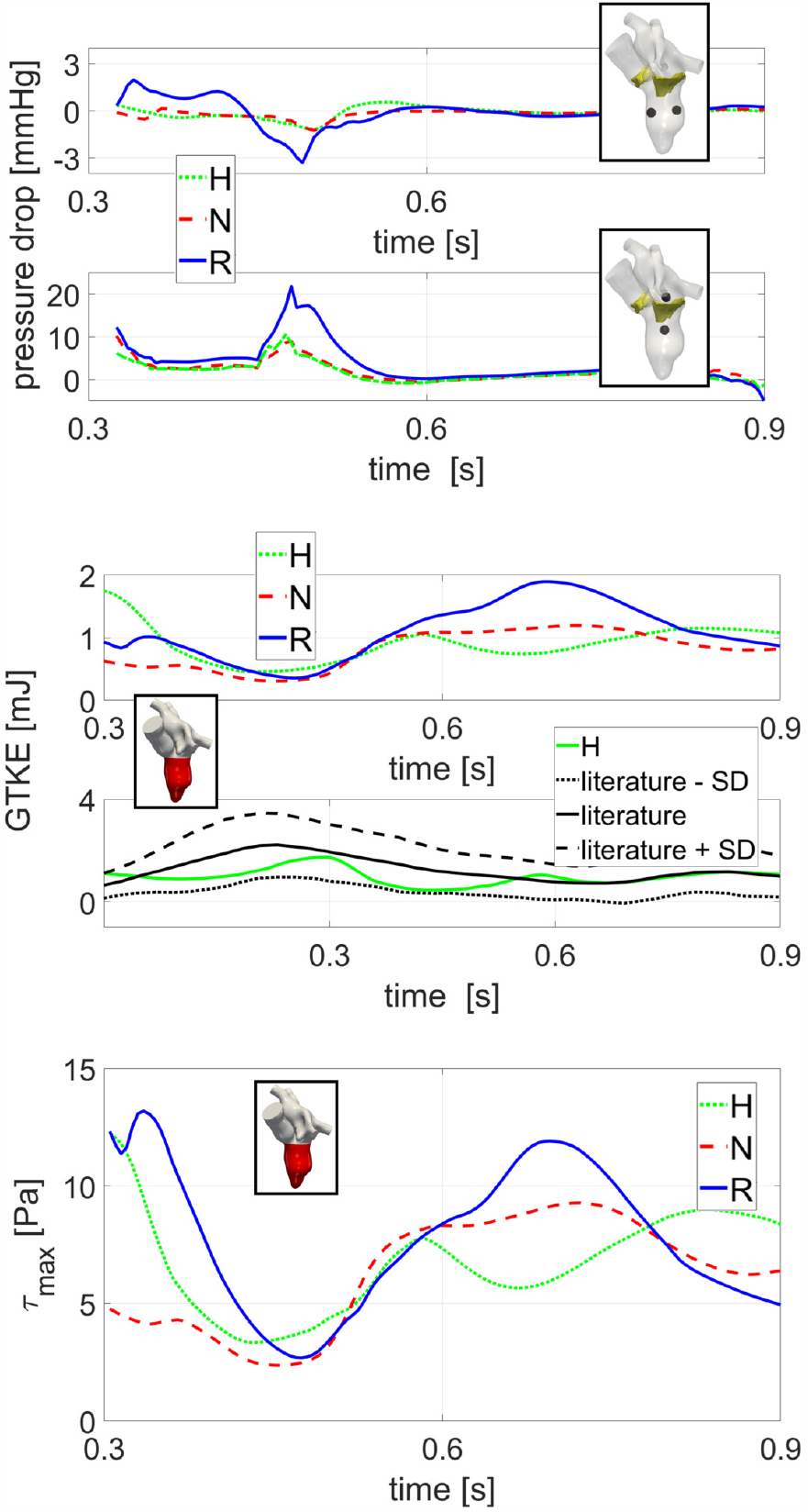
Top: Evolution in time, during diastole, of the pressure drops in the transversal and transvalvular directions; Middle: Evolution in time of GTKE integrated over the left ventricle volume during the diastolic phase (top) and comparison with respect to an average reference [64] (bottom); Bottom: Evolution in time of *τ*_*max*_ integrated over the left ventricle volume (bottom). Scenarios: Healthy (H), Neochordae (N), and Resection (R).

In Figure 9, we reported on a selected longitudinal slice the spatial distribution of the amount of vorticity along the direction perpendicular to the slice at the peak of E-wave t2 and at the peak of diastasis t3. Positive values of the vorticity represent a clockwise direction of the vortexes. In the same figure we also plotted TKE at the same time instants. We noticed that at t2 all the three scenarios exhibited a similar pattern with two vortex rings developing along the two mitral leaflets characterized by opposite directions. Moreover, the largest values of TKE were found right below the anterior leaflet. Furthermore, we observed the existence of turbulent regions in correspondence of the apex especially in R, attributed to the presence of pronounced eddies, see also Figure 6. At t3, we observe a significant, well-defined clockwise vortex in the center of the ventricle for H and N, whereas for R multiple disorganized vortexes were found. This led to high values of TKE for R, concentrated in the LVOT and in the central region of the ventricle. In Figure 8, middle, we reported the trend in time of the Global Turbulent Kinetic Energy (GTKE, i.e. TKE integrated over the ventricle) during the diastolic phase. Notice the large values featured by R especially during diastasis, whereas similar lower values were observed for H and N. This results are in accordance with the average values reported in Table 1. In the same figure, we reported also GTKE values obtained as the average among several healthy cases with 4D flow MRI [64], highlighting very similar values for what we found in the H scenario.

**Fig. 9.**
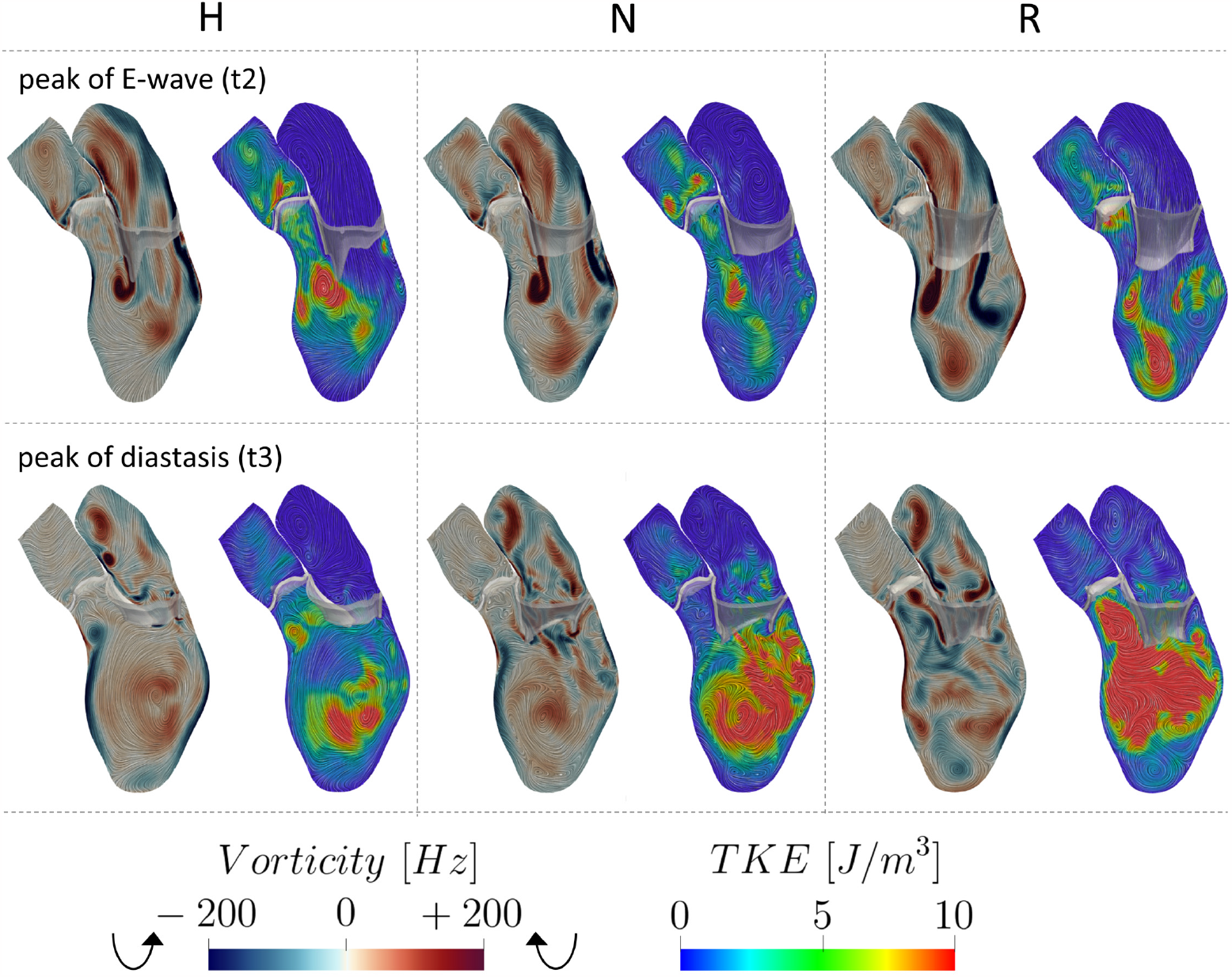
Spatial distribution of the vorticity and TKE at the peak of E-wave t2 and peak of diastasis t3, for the three scenarios Healthy (H), Neochordae (N) and Resection (R).

In Figure 10, we reported some quantities useful in view of a clinical analysis, such as the capability of washing out the ventricular apex and the hemolysis formation. Specifically, in the top figure we reported the spatial distribution of the Flow Stasis (FS) in the ventricular apex, where maximum values are attained. We can observe that R features lower values of FS than H and N cases. To quantify these differences we reported in Table 1 the percentage of volume of interest with FS greater than a representative threshold of 80%. This suggests a possible better ability of R to washout ventricular blood in the apex with respect to H and, especially, to N. Notice that, the analysis performed with other values of the threshold led to the same conclusions (percentage of area below the threshold lower in R).

**Fig. 10.**
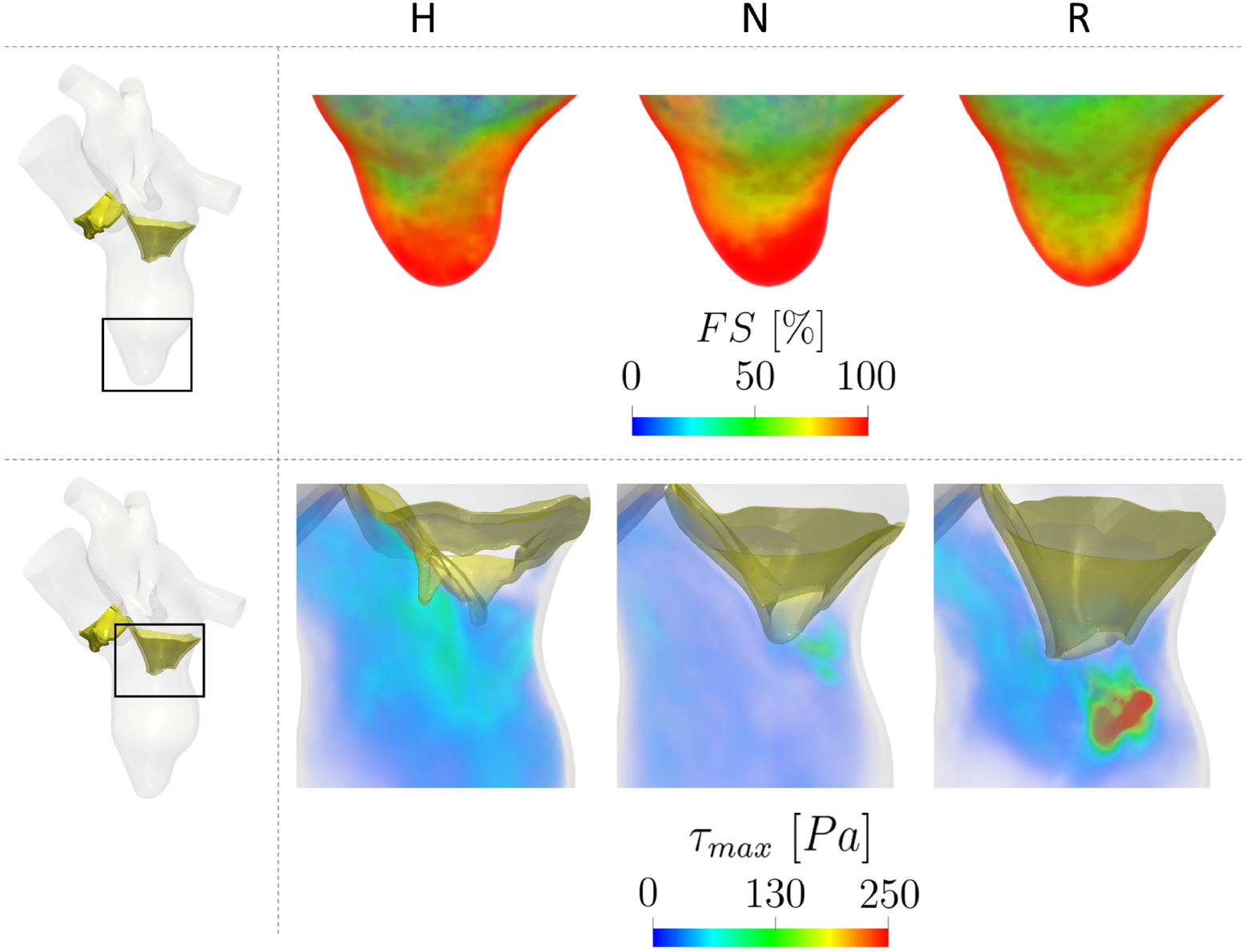
Top: Spatial distribution of FS(**x**) in correspondence of the ventricular apex; Botttom: Spatial distribution of *τ*_*max*_ in correspondence of the mitral valve during its opening. Scenarios: Healthy (H), Neochordae (N), and Resection (R).

In Figure 10, bottom, we reported the spatial distribution of the turbulent forces *τ*_*max*_ in a region of interest around the mitral valve during its opening. We noticed that, even if none of the three scenarios exceeded the critical threshold of 800 Pa [63], R gives rise to values up to 640 Pa (just below the mitral orifice), whereas H and N featured maximum values of 210 and 230 Pa, respectively. Accordingly, in Figure 8, bottom, we reported during the diastolic phase the evolution in time of the average *τ*_*max*_ within the ventricle. We notice that R featured larger values during the mitral opening than H and N, which displayed a similar trend. This is in accordance with the *τ*_*max*_ distribution at the valve opening reported in Figure 10, bottom. Also, the values of average-in-space *τ*_*max*_ are larger for R during diastasis, in correspondence of elevated values of GTKE, see Figure 8, middle. The time-average values of *τ*_*max*_ were reported in Table 1, confirming the significant attitude of R to develop large turbulent forces.

## 4 Discussion

In this work, we performed an image-based computational fluid dynamic study in the left heart to compare the haemodynamics in presence of two surgical reparative techniques (Neochordae and Resection) for the treatment of prolapse that may result in primary mitral regurgitation. To the best of the authors’ knowledge, this is the first computational study aiming to investigate the blood dynamics in presence of different mitral valve reparative techniques.

We found that the Resection technique reduces the mobility of the posterior (operated) leaflet, see Figures 1 and 5. This mainly influences the behaviour of the diastolic jet during the E-wave, which is directed in case of the Resection technique more towards the apex than in the other two cases, see Figure 6. On the contrary, the Neochordae technique features a diastolic jet more similar to the healthy case in terms of direction (developing more towards the ventricle wall) and vortexes formation, see Figures 6 and 9. The different direction of the jet allows blood in the Resection technique to reach the apex more quickly and with more pronounced washout compared to the other two cases. This is confirmed by the analysis performed on the Flow Stasis quantity (see Figure 10, top and Table 1), suggesting that the Resection technique may provide a greater protective role from potential thrombi formation [65].

The analysis of the pressure drops (Figure 8, top and Table 1) highlighted that within the ventricle the three scenarios exhibited comparable temporal evolutions (see Figure 8, top) and time-average values (see Table 1), and thus that the two operations lead to physiological values. As a consequence, according to the literature [30], the two techniques should not promote a non-physiological ventricular remodeling, characterized by a ventricle dilation which may occur in response to abnormal loading (pressure) conditions. From the analysis on the trans-valvular pressure drop we noticed that, although the Resection technique exhibited a time-average value of about 2 times greater than in the other cases (see Table 1), its trans-valvular pressure drop falls within the physiological range (0-5 mmHg) [66]. This suggests that both techniques ensure a proper mitral functioning during diastole.

The vorticity analysis highlighted that during the E-wave the Neochordae and Resection techniques showed a rotational behavior of the blood flow comparable to that of the healthy case. Specifically, all the scenarios are characterized by two different vortexes developing below the leaflets, see Figure 9, top. However, during diastasis, Resection technique amplifies the rotational dynamics of the blood and multiple and non-coherent eddies develop (see Figures 6 and 9, bottom). Instead, the Neochordae technique and the healthy scenario featured during diastasis the standard clockwise vortex in the middle of the ventricle, see Figure 9, bottom. The different behavior featured by the Resection technique at diastasis is mainly attributed to the reduced mobility of the posterior leaflet. This produces a jet which is directed (unlike the other two cases) towards the apex. As a consequence, according to the literature [29, 30], Resection technique may promote a non-physiological intracardiac vortex dynamics that may affect the heart efficiency, resulting in the worst scenario to heart failure.

We observed that there exists a relationship between areas with large vorticity and areas with high turbulence formation, see Figure 9. In particular, although the time evolution and average values of the three scenarios are comparable (see Figure 8, middle and Table 1), during diastasis there is much more turbulence formation in the Resection technique, due to the presence of several ventricular swirling structures, see also Figure 6. As a consequence, according to the literature [30], the presence of pronounced fluctuations in the Resection technique may contribute to increase the ventricular effort during the heartbeat.

According to the definition of *τ*_*max*_ (see Section 2.3), turbulence may create the conditions also for hemolysis development, a phenomenon related to the destruction of red blood cells due to fluid forces. According to [63], values of *τ*_*max*_ exceeding 800 Pa are identified as conditions that may induce hemolysis. Our findings reveal that the Resection technique significantly approaches this threshold during the mitral opening. This may indicate that a pronounced hypo-mobility of the posterior leaflet experienced by the Resection technique could result in the creation of turbulent forces capable of causing damage to red blood cells, thus promoting hemolysis which represents a potential cause of failure in mitral valve repair [67].

We point out that all the results in this study have been obtained under the assumption that both the repair techniques yield to the same ventricular displacement of the healthy subject. This assumption holds true for those post-operative scenarios that follow a pre-operative condition characterized by an almost physiological myocardial displacement. This is the case, for example of prolapse not resulting in regurgitation or if the operation occurs during the early stages of mitral regurgitation [68, 69]. Specifically, our two operated cases were characterized by no regurgitation of the prolapse.

Accordingly, the two operated mitral valves were virtually inserted and adapted in the left heart of the healthy subject by preserving the ratio between the area of the annulus and the area of the mitral orifice measured for the patients from imaging at the peak of the E-wave. We believe that this could be a well accepted strategy if one wants to compare the effect of only one change (in our case the mitral valve geometry) on the output of interest (see e.g. [49] for the case of mitral valve prolapse, [11] for different systolic anterior motion degrees, [23] for a structural analysis of the comparison of the two reparative techniques).

## Limitations

Some limitations characterized this work:

1. We considered only two operated patients and one healthy subject. This was a consequence of the fact that we used advanced (not daily available) images of the mitral valve in order to perform highly accurate DIB-CFD simulations. However we notice that this was not a statistical study, rather we wanted to describe the physical processes underlying blood dynamics of mitral repaired patients;
2. We did not include the papillary muscles in the ventricle geometry. This is a common choice in computational studies, adopted also in [11, 25, 70–72], due to the difficulty to reconstruct them from MRI images. Nevertheless, although our outcomes seems to be in accordance with previous studies, their influence on the quantities of interest could be relevant and it will be the subject of future studies;
3. We did not consider the chordae tendineae. This may be of particular relevance for the Neochordae technique. However, for this comparison study we believed that this common choice should not affect the qualitative conclusions on the two operated scenarios too much.
4. Our mesh did not include any boundary layer to better capture the blood dynamic behavior close to the myocardial wall. This should be considered in future studies. However, we noticed that our mesh resolution obtained after a refinement study was able to satisfy the Pope criterion [58].

## Acknowledgments

The authors acknowledge the CINECA award under the ISCRA C initiatives, for the availability of high-performance computing resources and support (IsCa8 DIB-CFD, P.I. Lorenzo Bennati, 2023).

## Declarations

### Ethical approval

Ethical review board approval and informed consent were obtained from all patients.

### Author contributions

Acquisition of the clinical data: GP

Methodology: LB, CV

Image Reconstruction and numerical simulations: LB

Conceptualization: LB, GP, VG, GBL, CV

Formal analysis and investigation: LB

Interpretation of the results: LB, CV

Writing - Original draft preparation: LB

Writing - Review and editing: CV, GP, VG, GBL

Supervision: CV, GBL

